# Abasic site ring opening and DNA-protein crosslink reversal by the SRAP protein YedK

**DOI:** 10.1101/2022.06.07.495154

**Authors:** Katherine A. Paulin, David Cortez, Brandt F. Eichman

## Abstract

Apuirinic/apyrimidinic (AP, or abasic) sites in DNA are one of the most common forms of DNA damage. AP sites are reactive and form crosslinks to both proteins and DNA, are prone to strand breakage, and inhibit DNA replication and transcription. The protein HMCES protects cells from strand breaks, inhibits mutagenic translesion synthesis, and participates in repair of interstrand DNA crosslinks derived from AP sites by forming a stable thiazolidine DNA-protein crosslink (DPC) to AP sites in single-stranded DNA (ssDNA). Despite the importance of HMCES to genome maintenance and the evolutionary conservation of its catalytic SRAP (SOS Response Associated Peptidase) domain, the enzymatic mechanisms of DPC formation and resolution are unknown. Using the bacterial homolog YedK, we show that the SRAP domain catalyzes conversion of the AP site to its reactive, ring-opened aldehyde form, and provide structural evidence for the Schiff base intermediate that forms prior to the more stable thiazolidine. We also report two new activities, whereby SRAP reacts with polyunsaturated aldehydes at DNA 3’-ends generated by bifunctional DNA glycosylases and catalyzes direct reversal of the DPC to regenerate the AP site, which provide insight into possible mechanisms by which HMCES DPCs are resolved in cells.

## Introduction

Apurinic/apyrimidinic (AP, or abasic) sites are one of the most ubiquitous DNA lesions. AP sites arise from either spontaneous or DNA glycosylase-catalyzed hydrolysis of the *N*-glycosidic bond that links the modified base to the deoxyribose (1,2). Their impact on cellular processes results in large part from their instability and reactivity. In solution, an AP site exists as an equilibrium between a predominant cyclic furanose as a mixture of α- and β-hemiacetals and a ring-opened aldehyde form, the latter constituting approximately 1% of the total (3–5). This electrophilic aldehyde can react with exocyclic groups of nucleobases on the complimentary strand to generate interstrand DNA crosslinks (ICLs) (6,7) and with primary amines in proteins to generate DNA-protein crosslinks (DPCs) (8,9). The ring opened aldehyde form is also susceptible to base-catalyzed β-elimination of the 3’ phosphoryl group, generating a singlestrand break (10). AP sites occurring in ssDNA, such as those encountered during replication, can lead to stalled replication forks by inhibiting replicative polymerases (2,11–13). Replication forks that encounter an AP site on the template strand can lead to a double-strand break (DSB) (2,12,14,15).

AP sites in double-stranded (ds)DNA are repaired by the base excision repair (BER) pathway, but the fate of AP sites in ssDNA is not as well understood. During replication, AP sites can be bypassed by error-prone translesion synthesis (TLS) polymerases (11,14,16,17). Recently, an alternative, higher fidelity pathway for repair of replication-associated AP sites was discovered that involves the protein HMCES (9,18,19). Cells lacking HMCES exhibit elevated levels and delayed repair of AP sites, as well as increased double-strand breaks and mutation frequency from TLS (18). Further supporting that HMCES responds to AP lesions, HMCES-deficient cells are hypersensitive to nuclear expression of APOBEC3A, which catalyzes deamination of cytosine to uracil in ssDNA that is converted to an AP site after removal by UNG (20,21).

HMCES forms DPCs with AP sites in ssDNA but not dsDNA (18), which led to a model in which the HMCES DPC protects AP sites from nuclease cleavage and mutagenic TLS polymerases (18,20). *In vitro*, both intact and proteolyzed HMCES DPCs are resistant to cleavage by AP endonuclease 1 (APE1) (18,22). Typically, proteins covalently conjugated to DNA occur either as deleterious lesions (23–25) or as a catalytic intermediate in DNA strand cleavage (lyase) reactions (26–28). By contrast, the HMCES DPC is highly stable and persists in cells on the order of hours, and has been shown to ultimately be resolved by a proteolytic-dependent mechanism under specific conditions (18,22). In *Xenopus* extracts, HMCES DPCs form as intermediates in AP-ICL repair upon NEIL3 unhooking of the AP-ICL, and are substrates for SPRTN protease, which generates a DNA-peptide crosslink (DpC) (29,30). However, the mechanism by which HMCES DPCs are resolved in mammalian cells remains to be determined.

HMCES contains a catalytic SRAP (SOS Response Associated Peptidase) domain that is conserved across all domains of life (18,22,31). HMCES SRAP is similar in both sequence (29% identity / 43% similarity) and structure to *E. coli* YedK, with the highest degree of conservation within the DNA binding channel and at the active site (22,32). An invariant cysteine at amino acid position 2 (Cys2) constitutes the extreme N-terminus after aminopeptidase removal of Met1 (18,33) and is required for DPC formation *in vivo* and *in vitro*. Crystal structures of HMCES SRAP and YedK crosslinked to AP-DNA revealed that Cys2 forms a highly stable thiazolidine linkage with the ring-opened aldehyde form of the AP site (32,34–36), which helped explain the persistence of HMCES DPCs in cells. The SRAP active site contains highly conserved glutamate, histidine, and asparagine residues that contact the crosslinked AP site (18,31,37). Mutation of these residues reduces crosslinking activity without disrupting DNA binding activity *in vitro* (18,22,33,35,36) and increases sensitivity to oxidative stress or ionizing radiation in cells (18,19,38).

Despite the importance of HMCES in repair of AP sites, the mechanisms of DPC formation and resolution, and the roles of active site residues in these processes, are unknown. Here, we perform a biochemical and crystallographic analysis of the various steps involved in catalysis of DPC formation, using YedK as a model system. Our data provide evidence for AP site ring opening and Schiff base formation, both of which are necessary precursors to thiazolidine formation. The active site glutamate is involved in both processes, and the histidine contributes to ring opening. We find that YedK forms DPCs to cleaved DNA 3’-ends generated by DNA lyases. We also show that YedK catalyzes DPC reversal to reform a free AP site on the order of several hours *in vitro*, which has implications for resolution of the HMCES DPC in cells.

## Results

### Glu105 and His160 enable acid-base catalysis of DPC formation

The Cys2-linked, ring-opened AP site is stabilized by highly-conserved histidine, glutamate, and asparagine residues (**Fig. 1A**). In YedK, Asp75 hydrogen bonds to the carbonyl oxygen and the backbone amide nitrogen of Cys2, His160 hydrogen bonds to the hydroxyl (O4’) of the ring-opened AP site, and Glu105 interacts with the thiazolidine ring and with His160 (32,34–36). In our crosslinked YedK structures, a second conformer of Glu105 was observed in which the carboxylate hydrogen bonds with the phosphate 3’ to the AP site, strongly implying that Glu105 exists at least transiently in a fully protonated state (22). Previous mutational analyses of SRAP active site residues involved only alanine substitution and were performed at a single time point (18,22,35). To gain a more detailed understanding of the roles of the SRAP active site residues, we performed a kinetic analysis of variants that altered their hydrogen bonding or ionization potential. We verified by mass spectrometry that these mutants all lack the N-terminal methionine (**Supporting Table S1**). Thus, the active site residues do not play a role in N-terminal methionine removal from the bacterial protein, contrary to a previous report using human HMCES (33).

**Figure 1.**
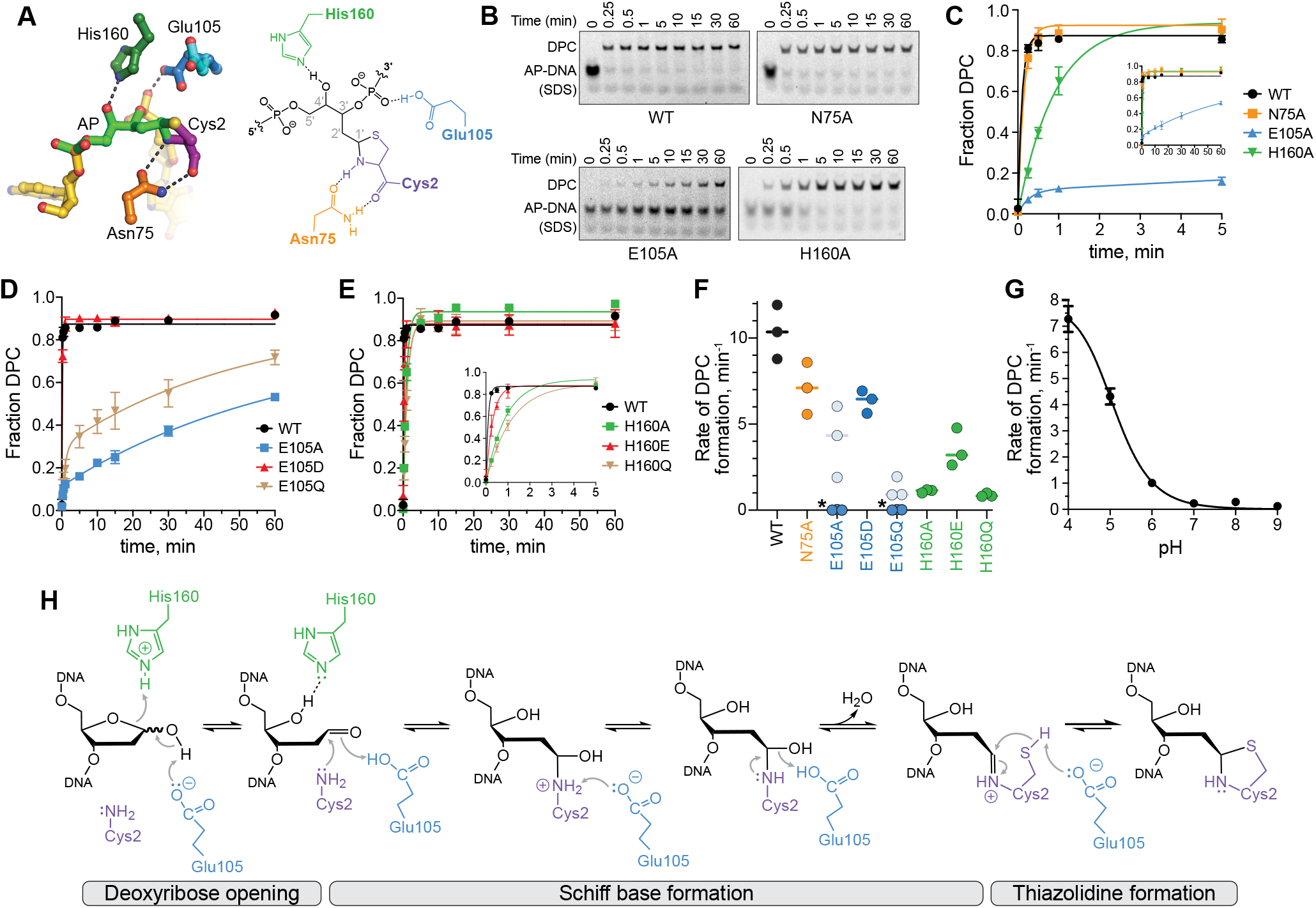
Glu105 and His160 enable acid-base catalysis of DPC formation. **A**. Active site of YedK DPC crystal structure (left) and schematic (right). **B**. Representative SDS-PAGE separation of uncrosslinked and crosslinked AP-DNA by wild-type and alanine mutant YedK. Crosslinking experiments were performed at 25 °C and pH 6. DNA bands were visualized with by FAM fluorescence. **C,D,E**. Kinetics of DPC formation of active site alanine mutants (C), Glu105 mutants (D), and His160 mutants (E) at 25 °C and pH 6 (mean ± SD, n=3). **F**. Rate constants derived from data in panels C-E. E105A and E105Q data were fit to a 2-phase exponential; *k*_fast_ is shown in light blue and *k*_slow_ is dark blue. Mean ± SD values are shown in Table S2. **G**. pH dependence of DPC formation for wild-type YedK at 18 °C (mean ± SD, n=3). Kinetic traces are shown in Fig. S1B. **H**. Proposed catalytic mechanism of YedK DPC formation.

Crosslinking kinetics were measured under single turnover conditions using a ssDNA oligo containing a centrally located AP site and a 5’-FAM label for visualization. In our assay, the rate of wild-type YedK DPC formation is a lower limit, as the reaction was nearly complete at our fastest time point. We first tested the kinetics of alanine point mutants (**Fig. 1B,C**). Surprisingly, Asn75, which was expected to position the N-terminal Cys2 for nucleophilic attack based on the structures, showed only a very modest decrease in crosslinking relative to wild-type YedK when mutated to alanine (**Fig. 1C,F**). In contrast, H160A exhibited at least 10-fold reduction in rate relative to wild-type. Alanine substitution of Glu105 had the largest effect of the three active site residues. The E105A crosslinking reaction was only 50% complete after 1 hour (**Fig. 1B-D**). These data are consistent with Glu105 and His160 as important for SRAP AP-site crosslinking, with Glu105 playing an essential role.

The proximity of Glu105 and His160 to the crosslink and to each other suggest that they participate in acid-base catalysis (34). We therefore examined the crosslinking kinetics of a E105Q and H160Q mutants, which cannot participate in acid-base chemistry but retain the same hydrogen bonding potential as the wild-type enzyme (**Fig. 1D-F, Supporting Figure S1A**). As with the alanine mutant, E105Q severely impacted YedK activity (**Fig. 1D,F; Table S2**), strongly suggesting that ionization of the carboxylate is important for DPC formation. Consistently, an E105D mutant only modestly impacted catalysis. The H160Q substitution also reduced the crosslinking rate 10-fold (similar to H160A), whereas an H160E mutant exhibited only a 3-fold reduction in crosslinking rate compared to wild type (**Fig. 1E,F; Table S2**). Thus, both Glu105 and His160 likely participate in acid-base catalysis rather than merely stabilize the substrate or transition state via hydrogen-bond stabilization. Consistent with this, YedK exhibits a strong pH dependence on the crosslinking rate with maximal activity at lower pH (**Fig. 1G; Fig. S1B**). The apparent midpoint of 5.1 in the pH profile is consistent with Glu105 acting as a general acid. The hydrophobic residues around the active site would raise the *pK_a_* of the glutamate, increasing the likelihood that it acts as both a proton donor and acceptor.

Based on the mutational data and the configuration of active site residues around the AP site, we propose the following catalytic mechanism for DPC formation in three main phases: AP site ring opening, Schiff base formation, and thiazolidine formation (**Fig. 1H**). In the first phase, both Glu105 and His160 likely catalyze ring opening of the AP deoxyribose ring from the furan to aldehyde form, whereby H160 acts as a general acid to protonate O4’ and Glu105 acts as a general base to deprotonate the hydroxide at C1’. In the second phase, Glu105 drives Schiff base formation by acting as both general acid and base to deprotonate Cys2 α-NH_2_ and to hydrolyze the hydroxyl at C1’. In the final step, Glu105 deprotonates the Cys2 sulfhydryl group necessary to close the thiazolidine ring.

### YedK catalyzes AP site ring-opening

In solution, the AP site is at equilibrium between a ring-closed 2’-deoxy-D-erythro-pentofuranose and a ring-opened aldehyde. The AP site exists primarily in the cyclic furanose form with only 1% of the sugar in the more reactive ring-opened aldehyde form (3,4). To investigate whether SRAP domains actively catalyze opening of the furan ring or simply capture a spontaneously formed aldehyde, we compared the rates of crosslink formation by two non-enzymatic probes to that of wild-type YedK under single-turnover conditions (**Fig. 2A-C; Fig. S2A-C**). The two non-enzymatic probes used were a YedK peptide consisting of the first 15 residues and including the N-terminal Cys2, and an aldehyde reactive probe, aoN-g, which reacts specifically to the aldehyde form of the AP site via an oxime linkage (39). The rates of YedK peptide and aoN-g probe crosslinking were 0.09 ± 0.005 min^-1^ and 0.03 ± 0.001 min^-1^, respectively, compared to 19.7 ± 2.4 min^-1^ for YedK. The 200-500-fold reduced rate in crosslinking by the two non-enzymatic probes suggests that YedK catalyzes AP site ring opening.

**Figure 2.**
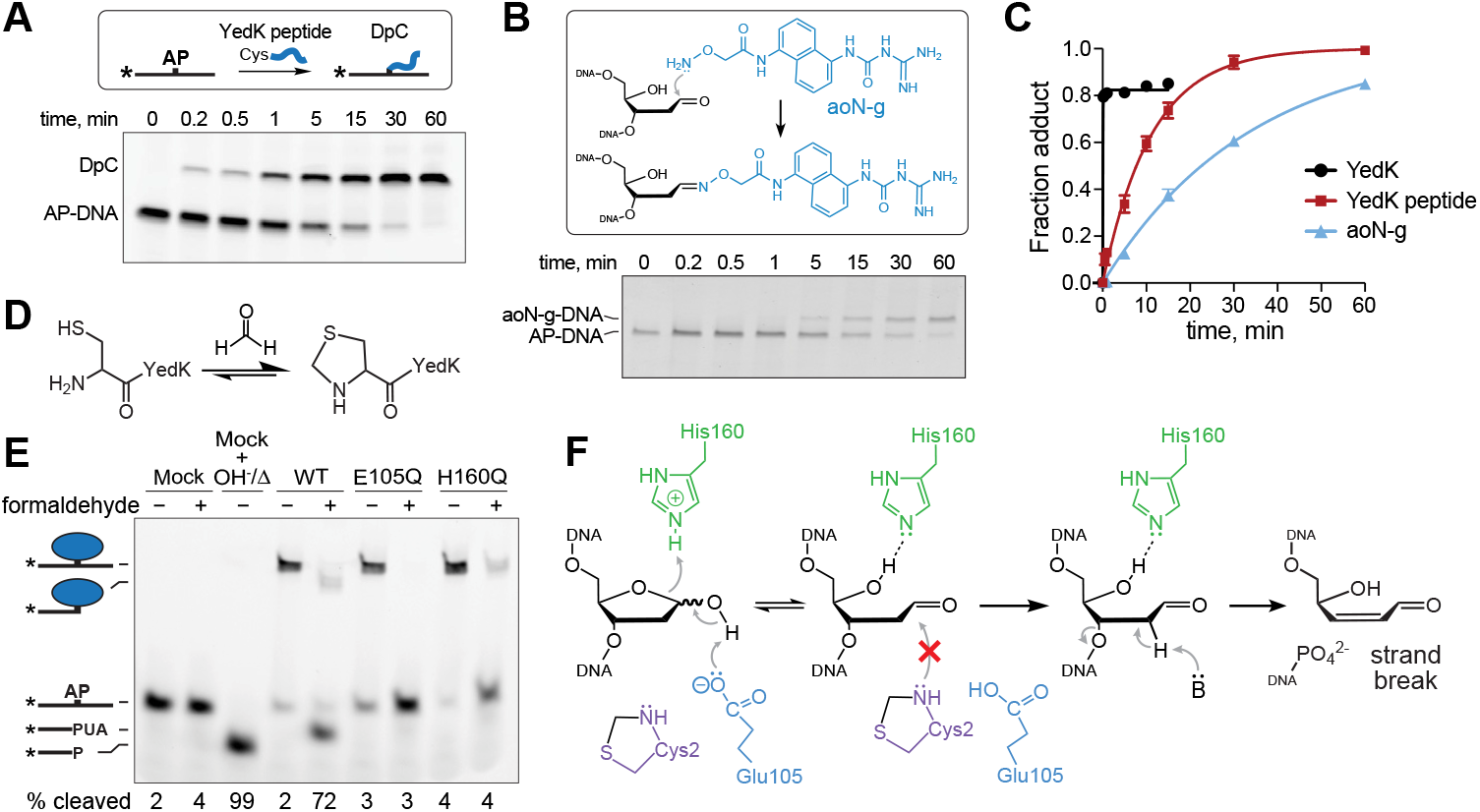
SRAP catalyzes AP site ring-opening. **A**. SDS-PAGE separation of AP-DNA crosslinked by YedK N-terminal peptide. **B**. Reaction of aldehyde reactive probe aoN-g to AP-DNA. **C**. Quantification of YedK peptide and aoN-g reaction with AP-DNA, compared to wildtype YedK (mean ± SD, n=3). **D**. Formaldehyde reacts with the YedK N-terminal Cys2 to form a thioazolidine. **E.** SDS PAGE of AP-DNA incubated with either buffer (mock) or native or formaldehyde-blocked YedK at 37 °C for 1 hour. PUA, 3’-phospho-α,β-unsaturated aldehyde (β-elimination product); P, 3’-phosphate (β,δ-elimination product). **F**. Blocking Cys2 with formaldehyde prevents YedK crosslinking to the ring-opened AP-site, leading to strand breakage.

To test this further, we selectively blocked the N-terminal Cys2 with formaldehyde, which reacts more efficiently with cysteine than other amino acids to form a thiazolidine ring (40–43) (**Fig. 2D**). Our proposed mechanism predicts that the α-NH_2_ group of Cys2 initiates DPC formation after the first step of AP site ring-opening (Fig. 1H). Thus, formaldehyde blocking of the N-terminus renders Cys2 unreactive toward the AP site, allowing us to examine the effects of Glu105 and His160 on the ring-opening step. As expected, blocking Cys2 in wild-type YedK inhibited DPC formation and led to strand cleavage (**Fig. 2E**), consistent with spontaneous β-elimination of the AP aldehyde previously observed with a C2A mutant (22) (**Fig. 2F**). In contrast, strand cleavage was not observed in Cys2-blocked E105Q and H160Q proteins, indicating that these residues are essential for formation of the reactive AP aldehyde. We verified that the loss of β-elimination in the formaldehyde-treated mutants was not the result of reduced DNA binding (**Fig. S2D**). Combined with the reduced rates of crosslinking by non-enzymatic probes, these data are consistent with Glu105- and His160-catalyzed AP ring opening by SRAP.

### Glu105 catalyzes formation of the Schiff base intermediate

SRAP DPC formation likely proceeds through a Schiff base intermediate formed by nucleophilic attack of C1’ of the AP site by the α-amino group of Cys2 (22,34,42,44). In the absence of the Cys2 thiolate side-chain, SRAP does not form DPCs (18,22,35) and instead generates DNA cleavage products indicative of DNA lyase activity (**Fig. 3A**). We previously provided evidence for the Schiff base intermediate by borohydride trapping of DPCs in YedK C2A and C2S mutants (22). To visualize this Schiff base intermediate, we determined a 1.8-Å crystal structure of YedK C2A with AP-DNA in the presence of borohydride. The electron density shows a linear linkage consistent with a reduced imine between the N-terminal alanine and the ring-opened AP site (**Fig. 3B**). This structure is highly similar to that of the wild-type YedK DPC (PDB ID 6NUA), with an RMSD of 0.71 Å for all Cα atoms. The DNA binding modality observed in the C2A DPC structure contains the same 90° kink and twist in the DNA backbone at the AP site observed in other SRAP-DNA structures (18,34–36) (**Fig. S3**).

**Figure 3.**
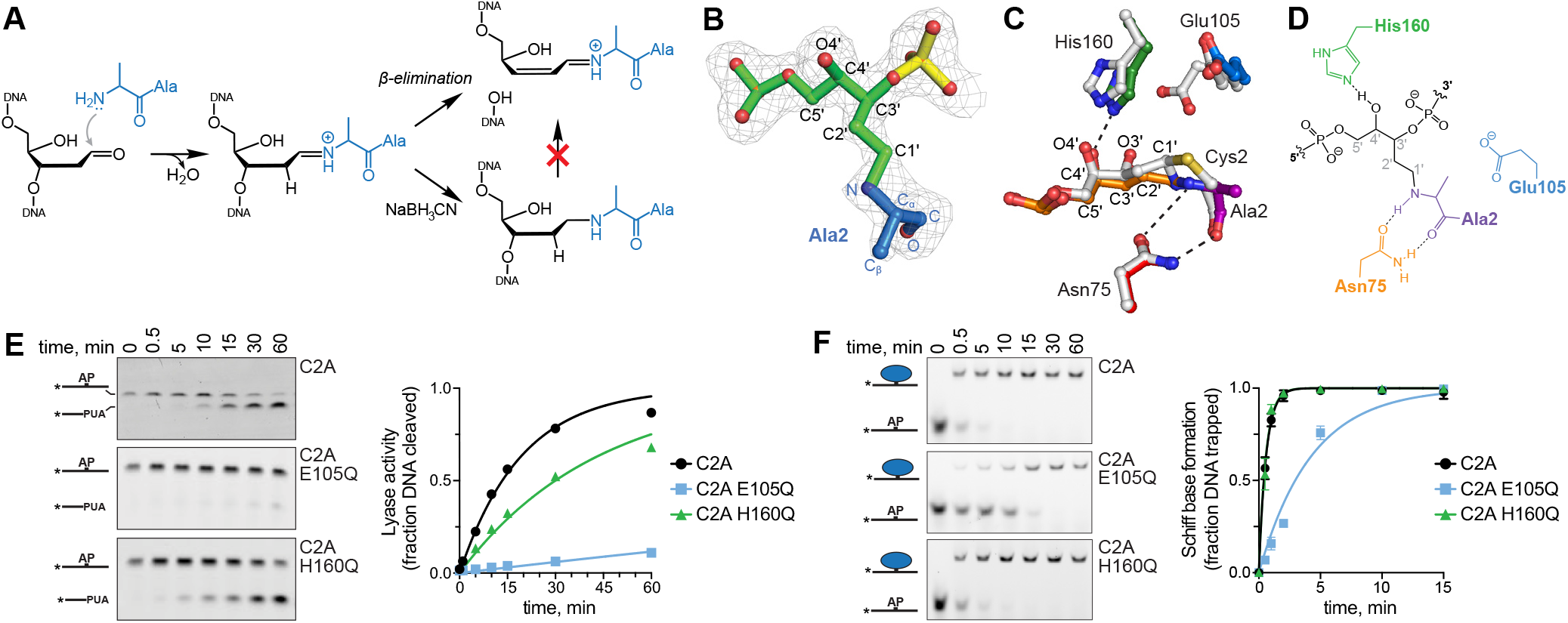
YedK DPC formation proceeds through a Schiff base intermediate. **A**. Borohydride reduction of the Schiff base formed between YedK C2A and AP-DNA prevents β-elimination. **B**. Crystal structure of the reduced YedK C2A DPC superimposed against 2F_o_ - F_c_ composite annealed omit electron density, contoured at 1σ. Ala2 is blue and the AP site is green. **C**. Superposition of wild-type (PDB ID 6NUA, white carbons) and C2A YedK (colored by residue) DPC structures. **D**. Schematic of the atomic interactions of the YedK C2A DPC. **E**. Kinetics of lyase activities of YedK C2A mutants in the absence of NaBH_3_CN. Quantitation of data from three independent experiments is shown on the right (mean ± SD, n=3). Rate constants derived from exponential fits to the data are 0.05 ± 0.002 (C2A), 0.002 ± 0.0001 (C2A E105Q), and 0.02 ± 0.001 (C2A H160Q). **F**. Kinetics of Schiff base formation between YedK C2A mutants and AP-DNA in the presence of NaBH_3_CN. Data from three experiments is quantified on the right (mean ± SD, n=3). Rate constants derived from exponential fits to the data are 1.7 ± 0.06 (C2A), 0.2 ± 0.02 (C2A E105Q), and 1.8 ± 0.09 (C2A H160Q).

The active site residues in the trapped Schiff base structure are positioned almost identically to those in the wild-type YedK DPC (**Fig. 3C,D**). The main notable difference in the C2A DPC structure is that Glu105 only exhibits one conformer; the interaction between carboxylate and DNA phosphate is not observed. In our proposed mechanism, Glu105 would catalyze Schiff base formation through deprotonation of Cys2 α-NH_2_ and hydrolysis of C1’. To investigate the roles of the active site residues E105 and H160 in catalyzing Schiff base formation, we determined the kinetics of both lyase activity and borohydride-trapped crosslinking of C2A E105Q and C2A H160Q double mutants (**Fig. 3E,F**). The C2A E105Q double mutant severely reduced both activities relative to C2A alone, whereas C2A H160Q had a lesser effect. The rates of lyase activity in C2A E105Q and C2A H160Q were 25-fold and 2.5-fold slower than C2A (**Fig. 3E**), further supporting an important role for Glu105 in the steps prior to Schiff base formation. Similarly, C2A E105Q reduced the rate of DPC formation in the presence of borohydride by 10-fold, whereas C2A H160Q showed the same rate as C2A (**Fig. 3F**). These data indicate that Glu105, but not His160, is important for Schiff base formation, consistent with our model (**Fig. 1H**).

### YedK reacts with AP lyase products

AP sites are susceptible to spontaneous and DNA lyase catalyzed strand cleavage through β-elimination of the 3’ phosphoryl group, which generates a single-strand break with a 3’-phospho-α,β-unsaturated aldehyde (3’-PUA) on one strand and a 5’-phosphate on the other (**Fig. 4A**) (10,45). The 3’-PUA may undergo further δ-elimination to liberate the ribose moiety, leaving a 3’-phosphate (3’-P). We tested the idea that SRAP could form a crosslink with the 3’-PUA by incubating AP-DNA with bifunctional DNA glycosylases Endonuclease III (Nth), Endonuclease VIII (Nei), or YedK C2A, all of which cleave AP-DNA (22,46,47), followed by incubation with wild-type YedK (**Fig. 4B**). In all cases, incubation with YedK resulted in the disappearance of the band corresponding to the lyase β-elimination product and the appearance of a corresponding DPC smaller in size to the YedK DPC formed with untreated AP-DNA. The amounts of the two DPCs in the three reactions were proportional to the amounts of uncleaved and cleaved AP-DNA from the lyase reaction, indicating that the lower molecular weight DPC is formed from the 3’-PUA. Consistent with the requirement for a reactive aldehyde, YedK did not react with the δ-elimination product of Nei. We also found that a preformed YedK DPC is refractory to DNA lyase cleavage by the glycosylases (**Fig. 4B**, right gel).

**Figure 4.**
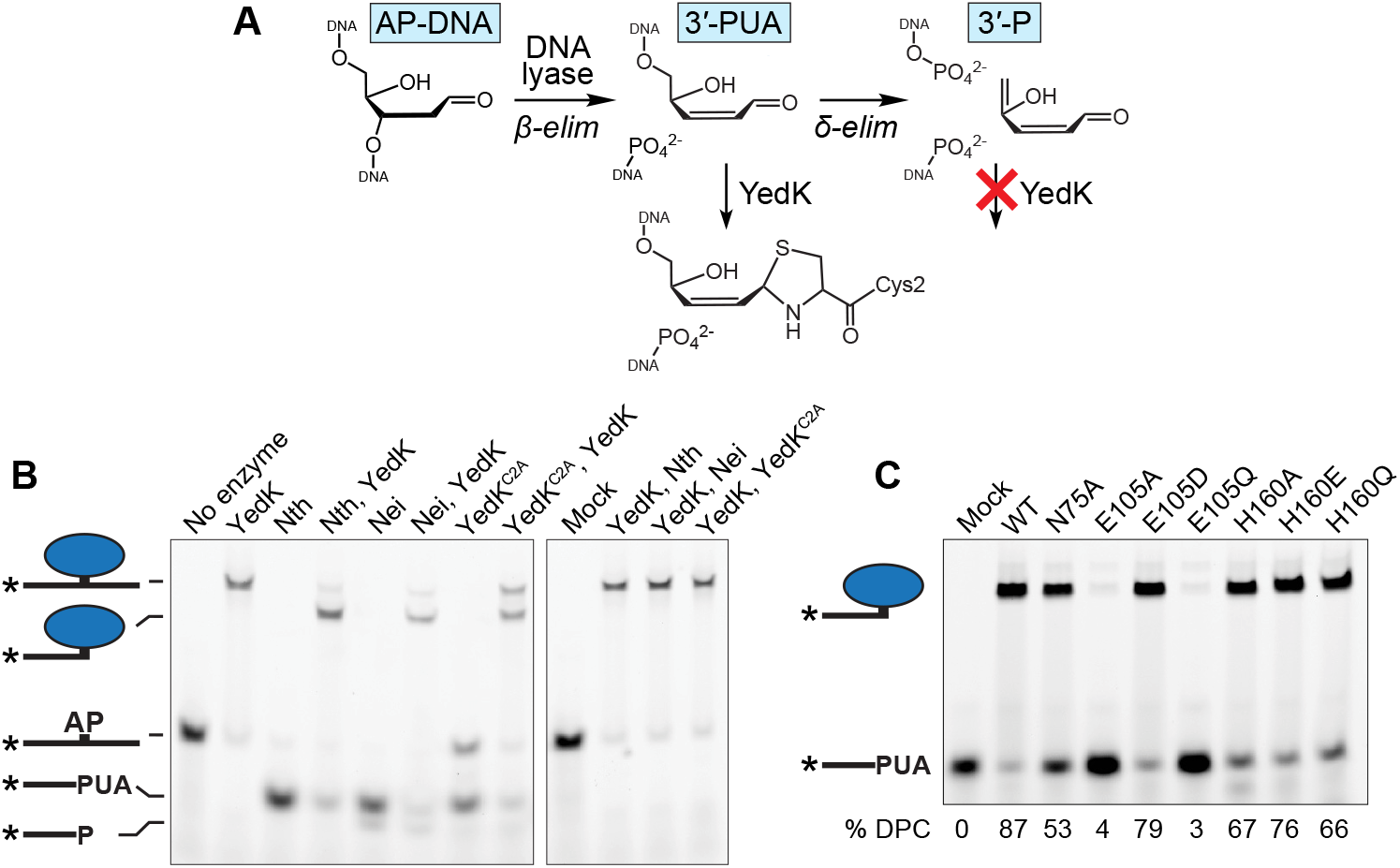
YedK reacts with AP lyase products. **A**. Reaction scheme for β-and δ-elimination by AP lyases. YedK can crosslink to the 3’-phospho-α,β-unsaturated aldehyde (3’-PUA) but not the 3’-phosphate (3’-P) of the δ-elimination product. **B**. YedK DPCs formed from the reaction products of AP lyases, Nth, Nei, and YedK C2A. Enzymes are listed in order of addition. **C**. Crosslinking of YedK mutants with the 3’-PUA formed by Nth. Reactions were carried out for 20 m at 37 °C.

We next tested our panel of active site mutants against 3’-PUA DNA substrates generated by Nth. Interestingly, N75A, which had only a modest effect on crosslinking to an internal AP site (Fig. 1C,F), was unable to fully crosslink the 3’-PUA after 20 minutes (**Fig. 4C**). Most notably, E105A and E105Q were refractory to 3’-PUA crosslinking, further supporting the role of Glu105 in formation of the Schiff base intermediate. Finally, the H160 mutants had a milder effect on 3’-PUA crosslinking relative to an internal site, consistent with this residue playing a role in ring-opening but not Schiff base formation.

### The SRAP DPC and DpC are reversible

SRAP DPCs are highly stable—on the order of hours in cells, and days *in vitro* at physiological temperature (18,22). The thiazolidine linkage itself is relatively stable and exists in an equilibrium with the Schiff base, with the thiazolidine favored by 5 orders of magnitude (42,43,48–50). We previously demonstrated thermal hydrolysis of the SRAP DPC after 10 minutes at 90 °C at pH 8.0 by following the protein component by SDS-PAGE (22). To elucidate the chemical nature of the DNA component of the hydrolysis product, we examined the sizes and reactivity of DNA products liberated from thermally denatured DPCs (**Fig. 5A**). Heating the DPC at 90 °C for 10 minutes at pH 6.5 completely hydrolyzed the DPC to generate two DNA products consistent with intact and nicked AP-DNA, both of which would contain a reactive aldehyde. The nicking observed is the result of spontaneous AP site hydrolysis (i.e., not YedK dependent) since the amount of nicked AP-DNA in the mock reaction is the same as the thermally denatured DPC reaction (Fig. 5A). Unlike free AP site, the DPC was refractory to base catalyzed β-elimination (**Fig. S4**). Addition of fresh YedK to the boiled DPC mixture generated two crosslinked species consistent with DPC formed from both DNA hydrolysis products. Thus, heat denaturation of the DPC leads to a direct reversal of the thiazolidine to regenerate a free, reactive AP site.

**Figure 5.**
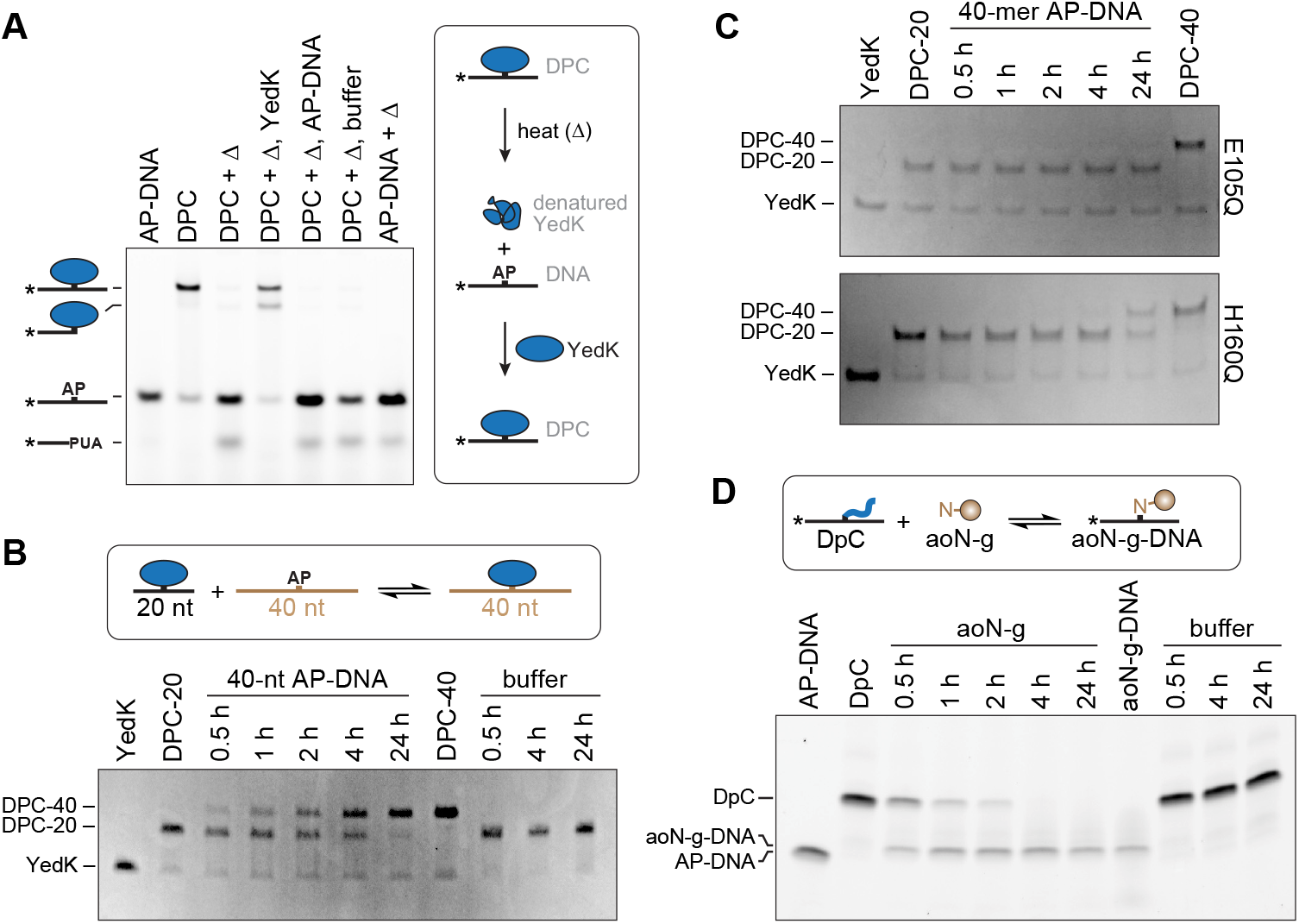
The SRAP DPC and DpC are reversible. **A**. Reaction of YedK DPC with heat (Δ), followed by either fresh YedK, AP-DNA, or buffer. An uncropped version of this gel is shown in Fig. S4. **B**,**C**. Time course of YedK DPC reversibility at 37°C for wild-type (B) and E105Q and H160Q mutants (C). DPC between YedK and 20-mer AP-DNA (DPC-20) was incubated with four-fold excess 40-mer AP-DNA or buffer and analyzed at the indicated time points. Free protein and DPC were visualized via Coomassie stained SDS-PAGE. **D**. YedK DpC pre-formed with 20-mer ssDNA was incubated with four-fold excess aoN-g or buffer. DNA bands were visualized with FAM fluorescence.

Since thiazolidine reversal and exchange with competing aldehydes is possible (51), we next tested whether the crosslink is reversible in solution under physiological conditions. DPC was pre-formed with a 20-mer oligodeoxynucleotide containing an AP site (DPC-20), followed by addition of 4-fold excess of 40-mer AP-oligodeoxynucleotide to trap any free AP-site liberated from a hydrolyzed DPC-20 (**Fig. 5B**). We observed the appearance of a 40-mer DPC (DPC-40) and a disappearance of DPC-20, consistent with direct reversal of the original DPC and reformation of DPC with the longer AP-containing oligo trap. The half-time of the exchange reaction under our experimental conditions was 2-4 hours. We verified that the reverse reaction was enzyme catalyzed, as DPC exchange was not observed after 24 hours with YedK E105Q and was severely slowed with H160Q (**Fig. 5C**). We also observed spontaneous reversal of a DpC formed with the YedK N-terminal peptide (**Fig. 5D**). In this case we used the aoN-g probe as a trap to capture any hydrolyzed DpC. As with the YedK DPC, we observed the disappearance of DpC and appearance of aoN-g-DNA over time, consistent with direct reversal of the DpC. Reversal of the DpC was about 2-4 fold faster than that of the DPC.

## Discussion

### Catalytic mechanism

Our data are consistent with Glu105- and His160-dependent, SRAP-catalyzed AP site ring opening to generate the reactive aldehyde necessary for attack by the Cys2 nucleophile. Three lines of evidence support this. First, wild-type YedK crosslinks to AP-DNA 2-3 orders of magnitude faster than the YedK peptide or the aldehyde reactive probe, aoN-g. These experiments were performed under saturating conditions to exclude diffusion rates from our interpretation. Secondly, by isolating the ring-opening step by formaldehyde blocking of Cys2 α-amino and thiolate side chain, we found that E105Q and H160Q mutants suppress spontaneous β-elimination of the AP site, suggesting that these residues are necessary to produce the reactive aldehyde form. Thirdly, our model predicts that His160 only plays a role in the ring-opening step. Consistently, YedK reactivity with a 3’-PUA, which effectively bypasses the requirement for the ring-opening step, were not as dependent on His160 as was the internal AP site crosslinking reaction.

Glu105 is by far the most important residue to DPC formation other than Cys2. In addition to facilitating ring-opening, likely by deprotonation of the hydroxide at C1’, Glu105 is also essential for Schiff base formation through its ability to deprotonate Cys2 α-NH_2_ and to protonate of the water leaving group from C1’. Our YedK C2A crystal structure under reducing conditions confirms that the DPC reaction proceeds through a Schiff base intermediate, and that the N-terminal amine is the initial crosslinking nucleophile. Interestingly, we did not observe a Glu105 conformer in contact with the DNA phosphate in the C2A DPC structure. This is consistent with our proposed mechanism in which Glu105 likely deprotonates the Cys2 sulfhydryl for thiazolidine ring closure in the last step of the reaction. In the absence of Cys2, Glu105 would remain in its anionic form.

Ionization of the Glu105 carboxylate is important for DPC formation since both E105A and E105Q had significant effects on catalysis. Both mutants exhibited biphasic kinetics at pH 6.0, with short burst (*k*_fast_) and longer slow (*k*_slow_) phases at least 2.5-fold and 1,000-fold slower than wild-type, respectively. The biphasic kinetics is not the result of reduced protein stability of the mutants since pre-incubation of protein before initiating the crosslinking reaction did not change the kinetics of E105A. This biphasic nature suggests that there are two forms of either the enzyme or the substrate at the onset of the reaction, one of which is primed for catalysis and bypasses the requirement for the enzyme in the initial step of the reaction. In the case of the enzyme, we speculate that two forms may exist that differ by the initial protonation state of the N-terminal α-amino group. Glu105 is positioned close enough to Cys2 to catalyze α-NH_3_^(+)^ deprotonation, which is required for Schiff base formation, and thus the initial burst may correspond to the population of the N-terminal amine already in the deprotonated state and the slow phase would correspond to the time required for spontaneous deprotonation. Alternatively, the initial burst may correspond to the small (1%) population of AP site that exists as the reactive aldehyde in solution, and the slow phase the result of the population of AP DNA in the more abundant, less reactive ring-closed state that requires Glu105 for activation (4,5).

### Reaction with DNA lyase products

HMCES reduces DNA double strand breaks (DSB) in cells, presumably by protection of the AP site from spontaneous or enzymatic cleavage at replication forks (18,20,22,29,36). Our finding that YedK is capable of crosslinking to DNA containing a 3’-PUA suggests additional roles for SRAP in DNA repair. The 3’-PUA DPC is consistent with reactivity of cysteine with α-β-polyunsaturated aldehydes (52). Specifically, we showed that YedK forms DPCs to the products of bacterial Nth and Nei glycosylases, raising the possibility that HMCES protects against the crosslinks to 3’-PUAs generated during abortive base excision repair, an activity that would contribute to HMCES-dependent reduction of spontaneous DNA strand breaks in cells. It is interesting to speculate that in addition to protecting AP sites from strand cleavage, the SRAP domain may also mark broken DNA ends for subsequent repair.

The 3’-PUA crosslinking activity raises the question of how SRAP recognizes a 3’-end generated by a DNA lyase. SRAP has a strong preference for AP sites in ssDNA (18,22,35) and can accommodate dsDNA on the 3’-side of the AP site (22,34). Lyase activity by a bifunctional DNA glycosylase would generate a nicked substrate with a duplex region 5’ to the reactive PUA, suggesting that fraying must occur for SRAP to gain access to the end. Interestingly, the ssDNA 3’ to the AP site, which would not be present in a 3’-PUA substrate, was disordered in several crystal structures of both HMCES SRAP domain and YedK (22,32,34,35), suggesting that the ssDNA is anchored to the protein mainly by the interactions 5’ to the AP site. Comparison of these structures to our YedK AP-DPC structure, in which the ssDNA containing an internal AP site fully extends across the active site, shows that the presence of DNA helps to orient the active site residues for catalysis (34). Binding to ssDNA containing a 3’-PUA would not be as stably bound to the protein and thus would require additional stabilization for catalysis. We speculate that this may be the role of Asn75, which is positioned to position Cys2 for nucleophilic attack on the AP site and was more important for YedK crosslinking to 3’-PUA than to an internal AP site.

### DPC reversibility

The mechanism by which the HMCES DPC is ultimately resolved is unknown. In human cells, resolution may take hours and involves proteasomal degradation (18). In *Xenopus* cell-free extracts the DPC is converted to a DNA-peptide crosslink (DpC) by SPRTN protease (29). How the DpC is resolved and whether the intact DPC is removed by alternative pathways are unclear. Here, we show that SRAP catalyzes direct reversal of its DPC back to a reactive AP site on a 2-4-hour timescale. We also observed a faster (1-2-hour) reversal of a DpC, indicating that reversal also occurs spontaneously from exposure of the thiazolidine to solvent. The difference in rates of the enzymatic and spontaneous crosslink reversal is consistent with thiazolidine hydrolysis occurring through the rate-limiting step of ring-opening to form a Schiff base (50), which is also susceptible to spontaneous hydrolysis. The relatively slow timescale of DPC reversal may be important to protect the AP-site during replication, which occurs over 7-8 hours (53). The 2-4-hour reversal of the DPC may allow for transient protection of the AP site until replication is completed in a specific region, at which point the DPC would be reversed, placing the AP site in the context of a ss/dsDNA junction for subsequent repair. Moreover, resolution of the DPC and DpC is likely cell-type or lesion-dependent (e.g., an AP site resulting NEIL3 unhooking of an AP-ICL versus depurination of a chemically modified nucleotide). The HMCES DPC is ubiquitylated in cells and this may target the protein for proteolysis or serve to recruit other DNA repair factors to the lesion (18,29). Additionally, there is evidence from *Xenopus* extracts that HMCES forms a DPC shortly after CMG helicase bypasses an unhooked AP-ICL, protecting the AP site from cleavage until TLS can proceed (29). Since HMCES reduces mutation frequency in U2OS cells (18,20) and TLS is an error-prone process, the delay of TLS by HMCES DPC likely allows for recruitment of an error-free bypass mechanism, such as template switching, in certain cell types. In B cells, DPC formation by HMCES reduces deletions during somatic hypermutation (SHM), and it has been proposed that subsequent TLS may be an outcome in SHM (54). Regardless of the mechanism, future investigation of SRAP DPC resolution should take into account the fact that the enzyme is capable of regenerating an AP site prior to the conclusion of DNA replication.

## Experimental procedures

### Site-Directed Mutagenesis

YedK mutants (C2A, C2A/E105Q, N75A, E105A, E105D, E105Q, H160A) were generated using the QuikChange Site-Directed Mutagenesis Kit (Agilent). The forward and reverse mutagenic extension reactions were performed separately to improve primer annealing, and the corresponding single stranded copies of the plasmid were combined. YedK point mutants (H160E, H160Q, C2A/H160Q) were generated using the Q5 Site-Directed Mutagenesis Kit (NEB). Mutant plasmids were sequence verified (GenHunter).

### Protein purification

*Escherichia coli* YedK and point mutants were expressed and purified as described (22). Briefly, His-tagged YedK was expressed in BL21 (DE3) cells at 16 °C for 16 h and purified by Ni-NTA affinity chromatography. The His-tag was cleaved and removed by a second Ni-NTA step. YedK was exchanged into S200 buffer (20 mM TRIS-HCl pH 8.0, 100 mM NaCl, 10% glycerol, 2 mM TCEP), concentrated, flash-frozen in liquid nitrogen, and stored at - 80 °C. Wild-type YedK used for biochemical experiments shown in figures 2A-B, 4B, and 5 was further purified by gel filtration on a 16/600 Superdex 200 column (GE Healthcare) in S200 buffer. The absence of the N-terminal methionine was verified via electrospray ionization (ESI) mass spectrometry (Vanderbilt Mass Spectrometry Core).

### Preparation of AP-DNA

AP-DNA was prepared by incubating 1 μM uracil-containing oligonucleotide with 0.6 U UDG (New England Biolabs) (55) in UDG Buffer (10 mM Tris-HCl pH 8.0, 50 mM NaCl, 10 mM MgCl_2_, 5 mM DTT) at 37 °C for 15 m. AP-DNA was prepared fresh for each reaction. Sequences of oligonucleotides used in biochemical assays are listed in Supporting Table S3.

### DNA-protein crosslinking kinetics

YedK DPCs were formed by incubation of 1 μM protein and 35 nM 5’-FAM-labeled AP oligonucleotide (FAM_U_Cy5) at 25 °C in Buffer A (20 mM Tris-HCl pH 6.0, 10 mM NaCl, 1 mM EDTA, and 5 mM DTT). Reactions were stopped at various time points by adding an equal volume of SDS Buffer (100 mM Tris-HCl pH 6.9, 16% glycerol, 3.2% SDS, 6% formamide, 0.5% β-mercaptoethanol) and incubating on ice. To confirm generation of AP sites, 35 nM 5’-FAM-labeled AP oligonucleotide in Buffer A was treated with 0.2 M NaOH for 3 m at 70°C. For Schiff base trapping, NaCNBH_3_ was added to the YedK and AP-DNA mixture to a final concentration of 50 mM. All samples were heated 70°C for 1 m prior to loading the gel. DPC and AP-DNA were separated on 4-12% Bis-Tris gels (Invitrogen) pre-run with MES SDS Running buffer (Invitrogen). 5’-FAM-labeled AP oligonucleotide was visualized on a Typhoon Trio (GE Healthcare) using excitation and emission wavelengths of 532 and 575 nm. Band intensities was quantified using GelAnalyzer 19.1 (www.gelanalyzer.com).

Experiments to measure C2A lyase kinetics were performed the same as DNA-protein crosslinking experiments, with C2A mutants used in place of wild-type YedK with the following modifications: reactions were incubated at 37 °C and stopped at given time points with equal volumes of Loading buffer (80% w/v formamide, 10 mM EDTA, 3 μg/μL Blue dextran) and reaction products were resolved via 10% polyacrylamide urea gels prerun in 0.5 X TBE buffer (50 mM Tris-HCl pH 8, 45 mM boric acid, 1 mM EDTA). Experiments to measure C2A Schiff base trapping kinetics were performed the same as DNA-protein crosslinking experiments, with C2A mutants used in place of wild-type YedK with the following modification: 50 mM NaBH_3_CN was added to the reaction mixture prior to taking the first time point.

For the pH dependence experiments, YedK DPCs were formed by incubation of 500 nM protein and 35 nM 5’-FAM-labeled AP oligonucleotide (FAM_U_Cy5) in Buffer B (20 mM Tris-HCl, 15 mM sodium citrate, 5 mM citric acid, 5 mM NaCl, 1 mM EDTA, 5 mM DTT, pH adjusted with HCl/NaOH and 0.22 μm filtered). AP-DNA was preincubated in reaction buffer at 18 °C for 10 m prior to addition of YedK and incubation at 18 °C at given time points over the course of 30 m. Reactions were quenched, analyzed by SDS-PAGE, and visualized by FAM fluorescence.

### Peptide and aoN-g crosslinking

Aldehyde reactive probe analog, aoN-g (39), was a gift from Yasuo Komatsu at National Institute of Advanced Industrial Science and Technology (AIST). YedK peptide consisting of the amino acids 2-16 (CGRFAQSQTREDYLA) was synthesized by Genscript. 50 nM 5’-FAM-labeled AP-DNA (FAM_U_20) was incubated at 25 °C with saturating concentrations of aoN-g (25 μM), YedK peptide (1 mM), or YedK (1 μM) in Buffer C (20 mM HEPES pH 7.0, 10 mM NaCl, 1 mM EDTA, 5 mM DTT). DNA-probe and DNA-peptide reactions were quenched by adding 8 μL reaction to 8 μL Stop Buffer (40 mM EDTA-Na2, 8M urea, 20 μM glutaraldehyde) and 4 μL of Loading buffer. YedK DPC reactions were stopped by adding an equal volume of SDS Buffer. All reactions were heated to 70 °C 1 m prior to loading gel. DNA-probe or DNA-peptide adducts were separated from AP-DNA on 15% polyacrylamide urea gels prerun in 0.5 X TBE buffer (50 mM Tris-HCl pH 8, 45 mM boric acid, 1 mM EDTA). DPCs were resolved via 4-12% Bis-Tris gels (Invitrogen) prerun in MES SDS running buffer (Invitrogen), and FAM-DNA visualized by fluorescence.

### N-terminal blocking

YedK and mutants stored in S200 buffer were thawed, spun 20,000 x g for 10 minutes, and diluted 10 μM in Buffer B at pH 7.0. Formaldehyde was added to a final percentage of 1% and reactions were incubated at 25 °C for 2 h before quenching by addition of 125 mM glycine. Reactions were buffer exchanged into fresh Buffer B pH 7.0 using G-25 desalting columns (Cytiva). 3.5 μM of formaldehyde-treated protein was incubated with 35 nM AP-DNA (FAM_U_35) in Buffer B pH 7.0 for 60 m at 37 °C. Reaction products were resolved via 4-12% Bis-Tris gels in MES running buffer and FAM-DNA visualized by fluorescence.

### DNA binding

Relative binding affinities of proteins in Fig. 2E were measured by fluorescence anisotropy using ssDNA (FAM_THF_15) containing a tetrahydrofuran (THF) abasic site analog. The THF strand contained 6-carboxfluorescein (FAM) at the 5’-end. Protein was titrated against 25 nM DNA in Buffer B in a 384-well plate for 20 min at 4°C. Fluorescence was measured using a BioTek Synergy H1 Hybrid Reader with a filter cube containing 485/20 nm excitation and 528/20 nm emission filters.

### X-ray crystallography

AP-DNA was prepared by incubating 50 μM 7-mer ssDNA [d(GTCUGGA]] with 10 U of uracil DNA glycosylase (UDG, New England Biolabs) in UDG Buffer at 37 °C for 1.5 h. YedK C2A protein was buffer exchanged into Buffer C. The Schiff base intermediate was trapped by incubating 24 μM YedK C2A with 25 μM AP-DNA at 25 °C for 5 m, adding NaBH_3_CN to a final concentration of 50 mM, and incubating at 25 °C for 18 h. DPC was further purified by gel filtration on a 16/300 Superdex 200 column (GE Healthcare). Fractions containing >90% DPC were pooled and buffer exchanged into 80 mM NaCl, 20 mM TRIS-HCl pH 8.0, 1 mM TCEP, 0.5 mM EDTA for crystallization experiments. DPC was crystallized by hanging drop vapor diffusion at 21 °C by mixing equal volumes of 2 mg/mL protein and reservoir solution containing 25% (wt/vol) PEG 3350 and 0.2 M NaH_2_PO_4_. Crystals were harvested 22 days after setting the drops, cryoprotected in 30% PEG 3350, 0.2 M NaH_2_PO_4_, and 10% (vol/vol) glycerol, and flash-cooled in liquid nitrogen.

X-ray diffraction data were collected at the Advanced Photon Source beamline 21-ID-F at Argonne National Laboratory and processed with HKL2000 (56). Data collection statistics are shown in Table S4. Phasing and refinement were carried out using the PHENIX suite of programs (57). Phases were determined by molecular replacement using the protein model from the YedK DPC structure (PDB accession 6NUA) followed by simulated annealing to eliminate model bias prior to further refinement. After refinement of atomic coordinates, temperature factors, and TLS-derived anisotropic B-factors, DNA was manually built in Coot, guided by 2m*E*_o_-D*E*_c_ and m*F*_o_-D*F*_c_ electron density maps. The Ala2-DNA crosslink as well as the entirety of the 7-mer ssDNA was readily apparent in the density maps. To minimize model bias, annealed m*F*_o_-D*F*_c_ omit maps were calculated by removing the Ala2 and the AP-site of the DNA. Geometry restraints for the linkage were generated from idealized coordinates of a reduced Schiff base (ChemDraw). The protein-DNA model was iteratively refined by energy minimization and visual inspection of the electron density maps. The final YedK-C2A-DNA model was validated using the wwPDB Validation Service and contained no residues in the disallowed regions of the Ramachandran plots. Refinement and validation statistics are presented in Table S4. All structural biology software was curated by SBGrid (58). Structure images were created in PyMOL (https://pymol.org). The structure was deposited in the Protein DataBank under accession code 8D2M.

### YedK reaction with lyase products

500 nM of YedK, Nth (EndoIII), Nei (EndoVIII), or YedK C2A was incubated with 35 nM 5’-FAM-labeled AP-DNA (FAM_U_35) in Buffer B pH 7.0 at 37 °C for 60 m. Aliquots were removed from each reaction, quenched by addition of an equal volume of SDS Buffer, and placed on ice. To each of the remaining reactions, fresh YedK or buffer as indicated was added to a final concentration of 500 nM and incubated at 37 °C for another 20 m. Reactions were stopped by adding 10 μL DPC reaction to 10 μL SDS Buffer and placed on ice. Reaction products were resolved on 4-12% Bis-Tris gels (Invitrogen) and FAM-DNA was visualized using excitation and emission wavelengths of 495 and 519 nm on a Chemidoc (BioRad).

### 3’-PUA reaction with YedK mutants

3’-PUA-DNA was generated by incubation of 1 μM 5’-FAM-labeled AP-DNA (FAM_U_35) with 20 U EndoIII/Nth (NEB) in Buffer B pH 7.0 at 37 °C for 60 m. 500 nM YedK was incubated with 35 nM 3’-PUA-DNA in Buffer B pH 7.0 at 37 °C for 60 m. Reactions were stopped with an equal volume of SDS Buffer. Reaction products were resolved on 4-12% Bis-Tris gels (Invitrogen) and visualized by FAM fluorescence.

### Crosslink reversal assays

For thermal DPC denaturation experiments, YedK DPC was formed by incubation of 35 nM 5’-FAM-labeled AP-DNA (FAM_U_35) with 500 nM YedK in Buffer C pH 6.5 for 60 m at 37 °C. The reaction was heated at 90 °C for 10 m to hydrolyze DPC, followed by addition of either buffer, fresh AP-DNA, or fresh YedK to the hydrolyzed DPC mixture and incubated for 10 m at 37 °C. Reactions were performed in HEPES rather than Tris buffer to avoid amines in the buffer being a confounding factor leading to strand cleavage (59). Reactions were stopped by adding an equal volume (10 μL) of SDS Buffer, products resolved on 4-12% Bis-Tris gels and visualized by FAM fluorescence.

Reversal trapping experiments were performed by incubating 10 μM 20-mer AP-DNA (FAM_U_20) and 2 μM YedK in Buffer C pH 7.0 for 18 h at 37 °C, which led to >90% DPC-20 formation. DPC-20 was incubated with a 4-fold excess (40 μM) of 40-mer AP-DNA (40_U) to trap any hydrolyzed DPC-20. Reactions were quenched with equal volumes of SDS Buffer. Each time point was initiated in reverse so that all reactions were quenched for the same length of time. Reaction products were resolved on 4-12% Bis-Tris gels and Coomassie stained for protein (22).

### DNA-peptide crosslink reversal over time

25 nM 20-mer AP-DNA (FAM_U_20) and 0.5 mM YedK peptide were incubated in Buffer C pH 7.0 for 1 h at 37 °C to form 100% DpC, after which 2 mM aoN-g probe was added and incubated at 37 °C for various times. Reactions were quenched by mixing 8 μL reaction, 8 μL 2X glutaraldehyde Stop Buffer, and 4 μL Loading buffer. Time points were initiated in reverse to maintain equal quenching times. Products were resolved on 15% polyacrylamide urea gels prerun in 0.5 X TBE buffer (50 mM Tris-HCl pH 8, 45 mM boric acid, 1 mM EDTA) and DNA was visualized via FAM fluorescence.

## Data availability

The model coordinates and structure factors for the crystal structure were deposited in the Protein DataBank under accession code 8D2M. All other data are included in the manuscript.

## Supporting information

This article contains four supporting figures and four supporting tables.

## Acknowledgements

The aoN-g probe was a gift from Yasuo Komatsu at National Institute of Advanced Industrial Science and Technology (AIST).

## Funding and additional information

This work was funded by grants from the National Institutes of Health (R35GM136401 to B.F.E., F31ES032334 to K.A.P, and R01ES030575 to D.C.). The content is solely the responsibility of the authors and does not necessarily represent the official views of the National Institutes of Health.

## Conflict of interest

The authors declare that they have no conflicts of interest with the contents of this article.

## Notes

### Competing Interest Statement

The authors have declared no competing interest.

## References

1. Lindahl, T., and Nyberg, B. (1972) Rate of depurination of native deoxyribonucleic acid. Biochemistry 11, 3610–3618

2. Krokan, H. E., and Bjørås, M. (2013) Base excision repair. Cold Spring Harbor perspectives in biology 5, a012583

3. Overend, W. G. (1950) 533. Deoxy-sugars. Part XIII. Some observations on the Feulgen nucleal reaction. Journal of the Chemical Society (Resumed), 2769–2774

4. Manoharan, M., Ransom, S. C., Mazumder, A., Gerlt, J. A., Wilde, J. A., Withka, J. A., and Bolton, P. H. (1988) The characterization of abasic sites in DNA heteroduplexes by site specific labeling with carbon-13. Journal of the American Chemical Society 110, 1620–1622

5. Wilde, J. A., Bolton, P. H., Mazumder, A., Manoharan, M., and Gerlt, J. A. (1989) Characterization of the equilibrating forms of the aldehydic abasic site in duplex DNA by oxygen-17 NMR. Journal of the American Chemical Society 111, 1894–1896

6. Yang, Z., Price, N. E., Johnson, K. M., Wang, Y., and Gates, K. S. (2017) Interstrand crosslinks arising from strand breaks at true abasic sites in duplex DNA. Nucleic acids research 45, 6275–6283

7. Price, N. E., Johnson, K. M., Wang, J., Fekry, M. I., Wang, Y., and Gates, K. S. (2014) Interstrand DNA-DNA cross-link formation between adenine residues and abasic sites in duplex DNA. Journal of the American Chemical Society 136, 3483–3490

8. Nakamura, J., and Nakamura, M. (2020) DNA-protein crosslink formation by endogenous aldehydes and AP sites. DNA repair, 102806

9. Thompson, P. S., and Cortez, D. (2020) New insights into abasic site repair and tolerance. DNA Repair (Amst) 90, 102866

10. Lhomme, J., Constant, J. F., and Demeunynck, M. (1999) Abasic DNA structure, reactivity, and recognition. Biopolymers: Original Research on Biomolecules 52, 65–83

11. Haracska, L., Unk, I., Johnson, R. E., Johansson, E., Burgers, P. M., Prakash, S., and Prakash, L. (2001) Roles of yeast DNA polymerases δ and ζ and of Rev1 in the bypass of abasic sites. Genes & development 15, 945–954

12. Hitomi, K., Iwai, S., and Tainer, J. A. (2007) The intricate structural chemistry of base excision repair machinery: implications for DNA damage recognition, removal, and repair. DNA repair 6, 410–428

13. Tsutakawa, S. E., Lafrance-Vanasse, J., and Tainer, J. A. (2014) The cutting edges in DNA repair, licensing, and fidelity: DNA and RNA repair nucleases sculpt DNA to measure twice, cut once. DNA repair 19, 95–107

14. Powers, K. T., and Washington, M. T. (2018) Eukaryotic translesion synthesis: Choosing the right tool for the job. DNA repair 71, 127–134

15. Fromme, J. C., and Verdine, G. L. (2004) Base Excision Repair. in Advances in Protein Chemistry, Academic Press. pp 1–41

16. Schaaper, R. M., Kunkel, T. A., and Loeb, L. A. (1983) Infidelity of DNA synthesis associated with bypass of apurinic sites. Proceedings of the National Academy of Sciences 80, 487–491

17. Andersen, P. L., Xu, F., and Xiao, W. (2008) Eukaryotic DNA damage tolerance and translesion synthesis through covalent modifications of PCNA. Cell research 18, 162

18. Mohni, K. N., Wessel, S. R., Zhao, R., Wojciechowski, A. C., Luzwick, J. W., Layden, H., Eichman, B. F., Thompson, P. S., Mehta, K. P., and Cortez, D. (2019) HMCES Maintains Genome Integrity by Shielding Abasic Sites in Single-Strand DNA. Cell 176, 144–153. e113

19. Srivastava, M., Su, D., Zhang, H., Chen, Z., Tang, M., Nie, L., and Chen, J. (2020) HMCES safeguards replication from oxidative stress and ensures error-free repair. EMBO Rep, e49123

20. Mehta, K. P. M., Lovejoy, C. A., Zhao, R., Heintzman, D. R., and Cortez, D. (2020) HMCES Maintains Replication Fork Progression and Prevents Double-Strand Breaks in Response to APOBEC Deamination and Abasic Site Formation. Cell Reports 31

21. Stenglein, M. D., Burns, M. B., Li, M., Lengyel, J., and Harris, R. S. (2010) APOBEC3 proteins mediate the clearance of foreign DNA from human cells. Nat Struct Mol Biol 17, 222–229

22. Thompson, P. S., Amidon, K. M., Mohni, K. N., Cortez, D., and Eichman, B. F. (2019) Protection of abasic sites during DNA replication by a stable thiazolidine protein-DNA crosslink. Nature Structural & Molecular Biology 26, 613–618

23. Covey, J. M., Jaxel, C., Kohn, K. W., and Pommier, Y. (1989) Protein-linked DNA strand breaks induced in mammalian cells by camptothecin, an inhibitor of topoisomerase I. Cancer Res 49, 5016–5022

24. Ide, H., Nakano, T., Salem, A. M. H., and Shoulkamy, M. I. (2018) DNA-protein cross-links: Formidable challenges to maintaining genome integrity. DNA Repair (Amst) 71, 190–197

25. Quiñones, J. L., Thapar, U., Wilson, S. H., Ramsden, D. A., and Demple, B. (2020) Oxidative DNA-protein Crosslinks Formed in Mammalian Cells by Abasic Site Lyases Involved in DNA Repair. DNA Repair, 102773

26. Zharkov, D. O., Rieger, R. A., Iden, C. R., and Grollman, A. P. (1997) NH2-terminal proline acts as a nucleophile in the glycosylase/AP-lyase reaction catalyzed by Escherichia coli formamidopyrimidine-DNA glycosylase (Fpg) protein. Journal of Biological Chemistry 272, 5335–5341

27. Gilboa, R., Zharkov, D. O., Golan, G., Fernandes, A. S., Gerchman, S. E., Matz, E., Kycia, J. H., Grollman, A. P., and Shoham, G. (2002) Structure of formamidopyrimidine-DNA glycosylase covalently complexed to DNA. Journal of Biological Chemistry 277, 19811–19816

28. Pommier, Y., Pourquier, P., Urasaki, Y., Wu, J., and Laco, G. S. (1999) Topoisomerase I inhibitors: selectivity and cellular resistance. Drug Resistance Updates 2, 307–318

29. Semlow, D. R., MacKrell, V. A., and Walter, J. C. (2022) The HMCES DNA-protein cross-link functions as an intermediate in DNA interstrand cross-link repair. Nat Struct Mol Biol

30. Wu, R. A., Semlow, D. R., Kamimae-Lanning, A. N., Kochenova, O. V., Chistol, G., Hodskinson, M. R., Amunugama, R., Sparks, J. L., Wang, M., Deng, L., Mimoso, C. A., Low, E., Patel, K. J., and Walter, J. C. (2019) TRAIP is a master regulator of DNA interstrand crosslink repair. Nature 567, 267–272

31. Aravind, L., Anand, S., and Iyer, L. M. (2013) Novel autoproteolytic and DNA-damage sensing components in the bacterial SOS response and oxidized methylcytosine-induced eukaryotic DNA demethylation systems. Biology direct 8, 20

32. Halabelian, L., Ravichandran, M., Li, Y., Zeng, H., Rao, A., Aravind, L., and Arrowsmith, C. H. (2019) Structural basis of HMCES interactions with abasic DNA and multivalent substrate recognition. Nature Structural & Molecular Biology, 1

33. Kweon, S.-M., Zhu, B., Chen, Y., Aravind, L., Xu, S.-Y., and Feldman, D. E. (2017) Erasure of Tet-Oxidized 5-Methylcytosine by a SRAP Nuclease. Cell reports 21, 482–494

34. Amidon, K. M., and Eichman, B. F. (2020) Structural biology of DNA abasic site protection by SRAP proteins. DNA Repair 94, 102903

35. Wang, N., Bao, H., Chen, L., Liu, Y., Li, Y., Wu, B., and Huang, H. (2019) Molecular basis of abasic site sensing in single-stranded DNA by the SRAP domain of E. coli yedK. Nucleic Acids Research 47, 10388–10399

36. Shukla, V., Halabelian, L., Balagere, S., Samaniego-Castruita, D., Feldman, D. E., Arrowsmith, C. H., Rao, A., and Aravind, L. (2019) HMCES Functions in the Alternative EndJoining Pathway of the DNA DSB Repair during Class Switch Recombination in B Cells. Molecular Cell 77, 384–394

37. Spruijt, C. G., Gnerlich, F., Smits, A. H., Pfaffeneder, T., Jansen, P. W., Bauer, C., Münzel, M., Wagner, M., Müller, M., and Khan, F. (2013) Dynamic readers for 5-(hydroxy) methylcytosine and its oxidized derivatives. Cell 152, 1146–1159

38. Pan, Y., Zuo, H., Wen, F., Huang, F., Zhu, Y., Cao, L., Sha, Q. Q., Li, Y., Zhang, H., Shi, M., Liang, C., Huang, J., Zou, L., Fan, H. Y., Ju, Z., Wang, H., and Shen, L. (2022) HMCES safeguards genome integrity and long-term self-renewal of hematopoietic stem cells during stress responses. Leukemia

39. Kojima, N., Takebayashi, T., Mikami, A., Ohtsuka, E., and Komatsu, Y. (2009) Construction of highly reactive probes for abasic site detection by introduction of an aromatic and a guanidine residue into an aminooxy group. J Am Chem Soc 131, 13208–13209

40. Kamps, J. J. A. G., Hopkinson, R. J., Schofield, C. J., and Claridge, T. D. W. (2019) How formaldehyde reacts with amino acids. Communications Chemistry 2, 126

41. Metz, B., Kersten, G. F., Hoogerhout, P., Brugghe, H. F., Timmermans, H. A., de Jong, A., Meiring, H., ten Hove, J., Hennink, W. E., Crommelin, D. J., and Jiskoot, W. (2004) Identification of formaldehyde-induced modifications in proteins: reactions with model peptides. J Biol Chem 279, 6235–6243

42. Kallen, R. G. (1971) Mechanism of reactions involving Schiff base intermediates. Thiazolidine formation from L-cysteine and formaldehyde. Journal of the American Chemical Society 93, 6236–6248

43. Ratner, S., and Clarke, H. T. (1937) The Action of Formaldehyde upon Cysteine. Journal of the American Chemical Society 59, 200–206

44. Just, G., Chung, B. Y., Kim, S., Rosebery, G., and Rossy, P. (1976) Reactions of oxygen and sulphur anions with oxazolidine and thiazolidine derivatives of 2-mesyloxymethylglyceraldehyde acetonide. Canadian Journal of Chemistry 54, 2089–2093

45. Purmal, A. A., Rabow, L. E., Lampman, G. W., Cunningham, R. P., and Kow, Y. W. (1996) A common mechanism of action for the N-glycosylase activity of DNA N-glycosylase/AP lyases from E. coli and T4. Mutat Res 364, 193–207

46. Melamede, R. J., Hatahet, Z., Kow, Y. W., Ide, H., and Wallace, S. S. (1994) Isolation and characterization of endonuclease VIII from Escherichia coli. Biochemistry 33, 1255–1264

47. Bailly, V., and Verly, W. G. (1987) Escherichia coli endonuclease III is not an endonuclease but a beta-elimination catalyst. Biochem J 242, 565–572

48. Schubert, M. P. (1936) COMPOUNDS OF THIOL ACIDS WITH ALDEHYDES. Journal of Biological Chemistry 114, 341–350

49. Cook, A. H., and Heilbron, I. M. (1949) THIAZOLIDINES. in Chemistry of Penicillin (Clarke, H. T., Johnson, J. R., and Robinson, R. eds.), Princeton University Press. pp 921–972

50. Canle, M., Lawley, A., McManus, E. C., and O’Ferrall, R. A. M. (1996) Rate and equilibrium constants for oxazolidine and thiazolidine ring-opening reactions. in Pure and Applied Chemistry

51. Saiz, C., Wipf, P., Manta, E., and Mahler, G. (2009) Reversible thiazolidine exchange: a new reaction suitable for dynamic combinatorial chemistry. Org Lett 11, 3170–3173

52. Esterbauer, H., Ertl, A., and Scholz, N. (1976) The reaction of cysteine with α,β-unsaturated aldehydes. Tetrahedron 32, 285–289

53. Masai, H., Matsumoto, S., You, Z., Yoshizawa-Sugata, N., and Oda, M. (2010) Eukaryotic chromosome DNA replication: where, when, and how? Annu Rev Biochem 79, 89–130

54. Wu, L., Shukla, V., Yadavalli, A. D., Dinesh, R. K., Xu, D., Rao, A., and Schatz, D. G. (2022) HMCES protects immunoglobulin genes specifically from deletions during somatic hypermutation. Genes Dev 36, 433–450

55. Krokan, H. E., Saetrom, P., Aas, P. A., Pettersen, H. S., Kavli, B., and Slupphaug, G. (2014) Error-free versus mutagenic processing of genomic uracil--relevance to cancer. DNA Repair (Amst) 19, 38–47

56. Otwinowski, Z., and Minor, W. (1997) Processing of X-ray diffraction data collected in oscillation mode. Methods Enzymol. 276, 307–326

57. Adams, P. D., Afonine, P. V., Bunkóczi, G., Chen, V. B., Davis, I. W., Echols, N., Headd, J. J., Hung, L.-W., Kapral, G. J., and Grosse-Kunstleve, R. W. (2010) PHENIX: a comprehensive Python-based system for macromolecular structure solution. Acta Crystallogr. D Biol. Crystallogr. 66, 213–221

58. Morin, A., Eisenbraun, B., Key, J., Sanschagrin, P. C., Timony, M. A., Ottaviano, M., and Sliz, P. (2013) Collaboration gets the most out of software. eLife 2, e01456

59. Haldar, T., Jha, J. S., Yang, Z., Nel, C., Housh, K., Cassidy, O. J., and Gates, K. S. (2022) Unexpected Complexity in the Products Arising from NaOH-, Heat-, Amine-, and Glycosylase-Induced Strand Cleavage at an Abasic Site in DNA. Chemical Research in Toxicology

