## Supplemental Figures and Tables for "Abasic site ring opening and DNA-protein crosslink reversal by the SRAP protein YedK"

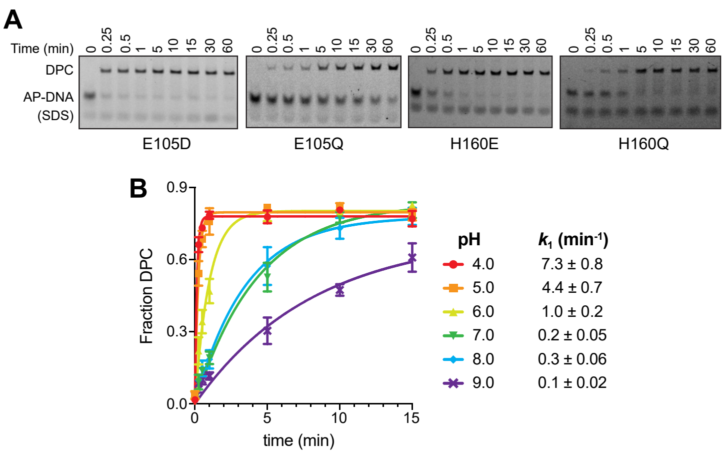


**Figure S1**. **A**. Representative denaturing PAGE gels of AP-DNA crosslinking by YedK point mutants. Bands were visualized by FAM fluorescence. **B**. Kinetics of crosslinking at different pH (mean ± SD, n=3). Reactions were performed at 18 °C. Rate constants derived from exponential fits to the curves are shown on the right and plotted in Fig. 1G.

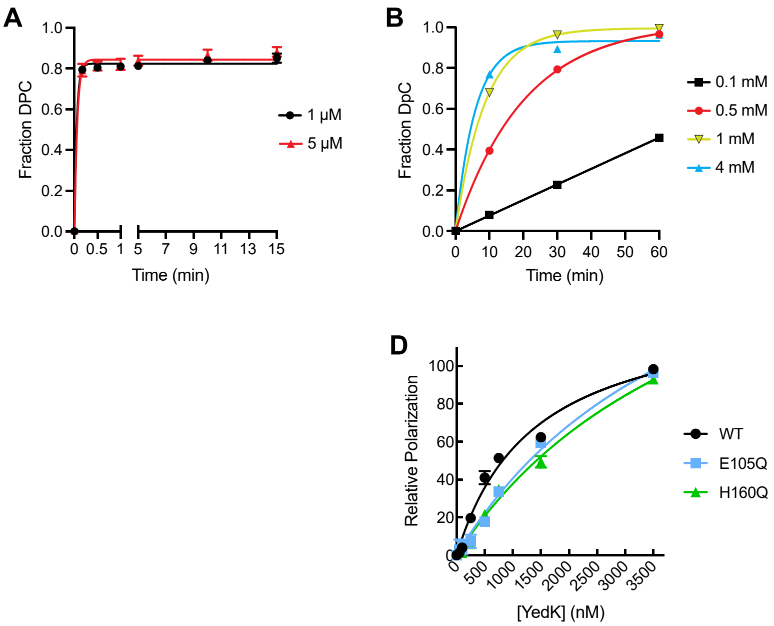


**Figure S2. A-C**. Determination of saturating conditions for crosslink formation using increasing concentrations of (A) YedK, (B) YedK peptide, and (C) aoN-g probe. **D**. Binding of formaldehyde-blocked YedK to ssDNA containing a 5′-FAM label and a centrally located tetrahydrofuran (THF) abasic site analog. Binding was monitored by a change in fluorescence polarization as protein was titrated against DNA. Dissociation constants ($K_d$) derived from non-linear least squares fit of the data are 1.4 μM ± 0.007 (WT), 4.5 ± 0.02 (E105Q), and 4.9 ± 0.02 (H160Q).

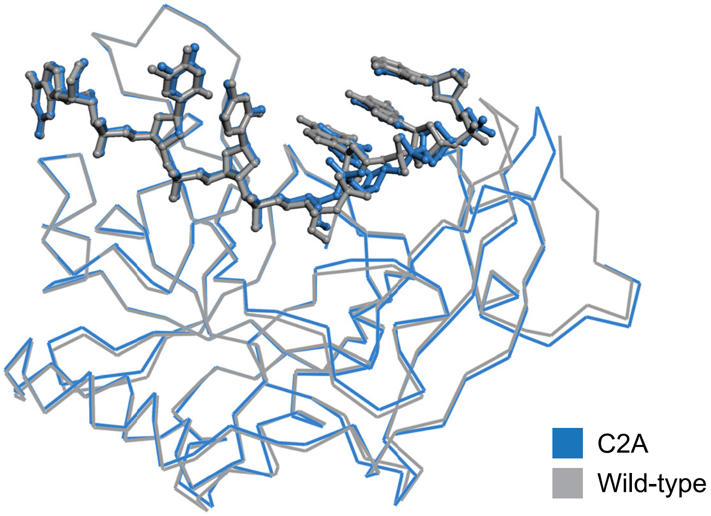


**Figure S3**. Superposition of the YedK C2A trapped Schiff base (this work) and the YedK DPC (PDB ID 6NUA) crystal structures.

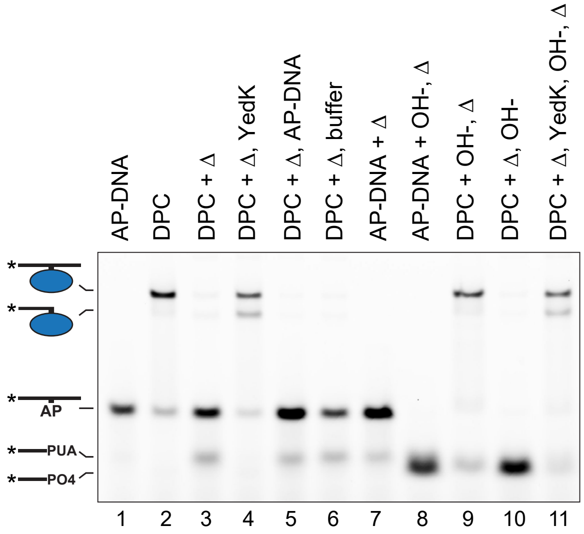


**Figure S4**. YedK DPC is refractory to strand breakage by alkaline pH. AP-DNA or DPC were reacted with hydroxide and heat (Δ) in the orders shown. Lanes 1-7 show that YedK DPC abolished after boiling can be re-formed with addition of fresh YedK but not DNA or buffer. Lanes 8-11 show that YedK DPC is refractory to cleavage by sodium hydroxide. DNA bands were visualized with FAM fluorescence. A cropped version of this gel containing lanes 1-7 is shown in Fig. 5A.

### Supporting Tables

**Table S1.** ESI-MS values to verify removal of N-terminal methionine

| YedK mutant | Calculated MW with Met1 | Calculated MW without Met1 | Observed MW |
|---|---|---|---|
| WT | 25709.14 | 25577.95 | 25572.5 |
| N75A | 25666.11 | 25534.92 | 25527.8 |
| E105A | 25651.10 | 25519.91 | 25511.9 |
| E105D | 25695.11 | 25576.96 | 25589.02 |
| E105Q | 25708.15 | 25576.96 | 25577.5 |
| H160A | 25643.08 | 25511.88 | 25529.6 |
| H160E | 25701.11 | 25569.92 | 25552.7 |
| H160Q | 25700.13 | 25568.93 | 25578.8 |
| C2A | 25677.08 | 25545.89 | 25540.1 |
| C2A/E105Q | 25676.09 | 25544.90 | 25520.8 |
| C2A/H160Q | 25668.07 | 25536.87 | 25527.8 |

**Table S2.** YedK mutant crosslinking rates [a]

| YedK mutant | $k_1$ (min$^{-1}$) | $k_2$ (min$^{-1}$) | Fold change (relative to WT) |
|---|---|---|---|
| WT [b] | ≥ 10.3 ± 1.6 | | 1 |
| N75A | 7.1 ± 1.5 | | 0.7 |
| E105A | 4.1 ± 2.1 | 0.01 ± 0.01 | 0.4 \| 0.001 |
| E105D | 6.3 ± 0.7 | | 0.6 |
| E105Q | 1.2 ± 0.7 | 0.02 ± 0.02 | 0.1 \| 0.002 |
| H160A | 1.1 ± 0.1 | | 0.1 |
| H160E | 3.5 ± 1.1 | | 0.3 |
| H160Q | 0.9 ± 0.1 | | 0.1 |

[a] Reactions were carried out at pH 6.0, 25 ºC. Values are mean ± SD (n=3)
[b] Value for wild-type is a lower limit

**Table S3.** Sequences of oligodeoxynucleotides used

| Name | Sequence (5'→3') | Figure(s) |
|---|---|---|
| FAM_U_Cy5 | FAM-d(CGGGCGGCGGCAUAGGGCGCGGGCCTTTTTT)-Cy5 | 1B-G, 3E-F |
| FAM_U_20 | FAM-d(TCTTCTGGTCUGGATGGTAGT) | 5B-D |
| 40_U | d(GGAATCTGACTCTTCTGGTCUGGATGGTAGTTAAGTCTTGT) | 5B-D |
| FAM_U_35 | FAM-d(ATGACTCTTCTGGTCUGGATGGTAGTTAAGTCTTGT) | 2A-C, 2E, 4A, 4C, 5A, 5D |
| FAM_THF_15 | FAM-d(TCTGGTC[THF]GGATGGT) | S2D |
| H36 | d(GTCUGGA) | 3B-C |

**Table S4.** X-ray data collection and refinement statistics for YedK C2A DPC [a]

| | |
|---|---|
| **Data collection** | |
| Space group | $P2_1$ |
| Cell dimensions | |
| *a*, *b*, *c* (Å) | 60.94 41.47 82.58 |
| *α*, *β*, *γ* (°) | 90.00, 95.65, 90.00 |
| Resolution (Å) | 50.00 - 1.83 (1.90 - 1.83) |
| $R_{sym}$ | 0.076 (0.667) |
| $R_{meas}$ | 0.101 (0.891) |
| Avg. *I*/σ*I* | 11.9 (1.4) |
| Completeness (%) | 99.53 (96.06) |
| Redundancy | 2.1 (2.1) |
| Wilson *B*-factor (Å$^2$) | 19.8 |
| **Refinement** | |
| Resolution (Å) | 37.03 - 1.82 (1.87 - 1.82) |
| No. reflections | 37,024 (3,583) |
| $R_{work}$ | 0.178 (0.254) |
| $R_{free}$[b] | 0.226 (0.307) |
| No. atoms[c] | |
| Protein | 3,500 |
| DNA | 246 |
| Water | 260 |
| Other | 0 |
| Avg. *B*-factors[c,d] (Å$^2$) | |
| Protein | 26.6 |
| DNA | 29.4 |
| Water | 27.8 |
| Other | - |
| R.m.s. deviations | |
| Bond lengths (Å) | 0.011 |
| Bond angles (°) | 1.109 |

[a] Statistics for the highest resolution shell are shown in parentheses.
[b] $R_{free}$ was determined from the 5% of reflections excluded from refinement.
[c] Riding hydrogen atoms were not included in no. atoms or avg. *B*-factors.
[d] Equivalent isotropic *B*-factors were calculated in conjunction with TLS-derived anisotropic *B*-factors
